# LC3-associated phagocytosis is impaired in monocyte-derived macrophages from systemic sclerosis patients

**DOI:** 10.1101/2024.03.21.586112

**Authors:** Quentin Frenger, Julie Lucas, Arthur Petitdemange, Luisa Path, Nadège Wadier, Sabine Depauw, Stéphane Giorgiutti, Yazhuo Gong, Hélène Merlin, Aurore Meyer, Mathilde Herber, Léa Jaquel, Philippe Mertz, Renaud Felten, Emmanuel Chatelus, Yannick Dieudonne, Aurélien Guffroy, Laurent Arnaud, Vincent Poindron, Jacques-Eric Gottenberg, Jean Sibilia, Anne-Sophie Korganow, Thierry Martin, Frédéric Gros

## Abstract

Autophagy is a fundamental catabolic process performed by a network of autophagy related (ATG) proteins. Some ATG proteins coordinate parallel roles in so-called “noncanonical” autophagy such as LC3-associated phagocytosis (LAP). Both autophagy and LAP share key functions in immunity and inflammation and have been linked to autoimmune diseases. Systemic sclerosis (SSc) is an autoimmune disease of unknown etiology characterized by excessive fibrosis in skin and multiple internal organs linked with an aberrant immune activation. Several polymorphisms of genes coding for ATG proteins, particularly in *ATG5*, are more frequent in SSc patients. We hypothesized that autophagy and/or LAP could be dysregulated in immune cells from SSc patients. No defect of canonical autophagy was found in lymphocytes and monocytes isolated from peripheral blood mononuclear cells of SSc patients. We then generated monocyte-derived macrophages and performed phagocytosis assays to assess LAP activity. While M0 macrophage polarization appears similar than in healthy donors, we showed that LAP is downregulated in SSc patients. We now need to understand the molecular mechanisms underlying LAP dysregulations. Future investigations leading to the discovery of LAP modulating drugs could then open new therapeutic options for SSc treatment.

**Key messages:**

- Polymorphisms of autophagy-related genes are associated with several autoimmune and autoinflammatory diseases, including SSc and SLE
- While autophagy has been shown to be dysregulated in circulating cells from SLE patients, no information is available for SSc
- We show here that autophagy is comparable between PBMCs from patients and matched controls
- We find a strong impartment of LAP, another ATG-dependent mechanism, in monocyte-derived macrophages from SSc patients
- As LAP is involved in efferocytosis and the regulation of inflammation, we propose that restoring LAP activity could be a therapeutic option to limit fibrosis and inflammation

## Introduction

Systemic sclerosis (SSc) is an autoimmune disease with fibrosis affecting skin and internal organs^1^. Dysregulation of fibroblast activity is linked to inflammation induced by both innate and adaptive immune cells^2^. Defects in the phagocytic compartment have been proposed to contribute to the disease as exemplified by unbalanced M1/M2 macrophage polarization observed in several studies^3^. Efferocytic defects that could be linked to biased macrophage regulation have been recently highlighted: monocyte-derived macrophages from SSc patients showed lower levels of SR-B1s scavenger receptor compared to healthy subjects leading to decreased uptake of apoptotic cells^4^. The same group showed that crystalline silica, a known environmental factor predisposing for SSc leads to efferocytosis impairment, associated with inhibition of M2 polarization and increased activity of the essential phagocytosis RhoA GTPase^5^. Interestingly inhibiting RhoA/ROCK pathway restores efferocytic capacity.

ATG (autophagy-related) proteins are molecular executors of macroautophagy (further mentioned as autophagy) but also other vesicular trafficking events. In adaptive immune cells, autophagy is essential for the survival of long-lived cells like memory lymphocytes and plasma cells^6^. In phagocytes, autophagy participates in the elimination of intracellular organisms, in the regulation of inflammation through limitation of inflammasome activity and type I IFN production, and promotes antigen presentation of cytosolic molecules^7^. A mechanism of intracellular vesicle decoration with LC3 (light chain 3, abbreviation of MAP1LC3B microtubules associated light chain 3, ortholog of yeast Atg8) in processes called “Atg8ylation” also plays specialized roles especially in phagocytes^8^. One form of Atg8ylation in macrophages is LC3-associated phagocytosis (LAP), the most characterized form of CASM (conjugation of ATG proteins to single membranes)^9,10^. Its integrated physiological roles are still unclear but LAP impacts efferocytosis, antigen presentation, and internalization vesicles trafficking. For example LC3 is necessary to address endocytic vesicles to TLR-containing compartments thus enhancing inflammation^11^. On the contrary, LAP activity limits in other settings IFN-I and IL-1β production^12^ during efferocytosis. Altogether, these findings suggest that ATG proteins could be central to chronic inflammatory disease pathogenesis.

Polymorphisms on *ATG5* has been associated with the onset of several inflammatory pathologies including systemic sclerosis^13^. Few studies addressed the role of ATG proteins in SSc, and none in immune cells from patients, especially phagocytes. As deregulations of autophagy and/or LAP could indeed contribute to pathologies like Crohn’s disease, systemic lupus erythematosus (SLE), multiple sclerosis, Sjögren disease or rheumatoid arthritis^14^, we propose that it could also contribute to SSc pathogenesis.

## Material and methods

### Patients and healthy donors

The patients included in this study suffer from dcSSc and are aged over 18 years. They were recruited from the clinic immunology department at “Nouvel Hôpital Civil” and Rheumatology department of “Hôpital de Hautepierre”, of Strasbourg. Cognate controls were purchased from the “Etablissement Francais du Sang” in Strasbourg. Patients fulfilled the American College of Rheumatology criteria for their respective diseases (Tables 1 and 2). They signed a consent form after being informed about the protocol and the aim of this study. Patients and controls are age and sex-matched and blood samples were taken in EDTA tubes and treated in the shortest delays.

**Table 1:** Systemic sclerosis patients informations. Age, sex, date of prelevement, age of first sign, clinical manifestations, ACR/EULAR scores, treatments at the time of prelevement and presence of auto antibodies. MMF : mycophenolate mofetil.

| Age | Sex | Prelevement date | Age of first sign | Clinical manifestations | ACR/EULAR score | Treatments | Auto-antibodies |
| --- | --- | --- | --- | --- | --- | --- | --- |
| 49 | M | 2019_01_22 | 48 | Lungs | 10 | None | SCL70 |
| 51 | M | 2021_06_07 | 48 | Skin, Lungs, Kidneys, Digestive | 21 | Cyclophosphamide | SCL70 |
| 78 | F | 2019_01_30 | 73 | Skin | 23 | MMT | RNA-Pol-III |
| 62 | F | 2019_02_08 | 51 | Skin, Heart, Lungs | 26 | Steroids >12,5mg | Centromere |
| 64 | F | 2021_05_04 | 51 | Skin, Heart, Digestive | 24 | N-C | Centromere |
| 62 | F | 2019_02_20 | 61 | Skin, Heart, Lungs | 23 | MMT, Steroids >12,5mg | SCL70 |
| 64 | F | 2019_05_06 | 58 | Skin, Heart, Kidneys | 16 | MMT, Steroids | nuclear only |
| 51 | F | 2019_03_14 | 50 | Skin | 19 | None | SCL70 |
| 44 | F | 2019_03_20 | 41 | Skin, Lungs | 21 | None | SCL70 |
| 38 | M | 2019_03_22 | 37 | Articular | 10 | None | SCL70 |
| 40 | M | 2021_03_21 | 37 | Articular | 10 | Methotrexate | SCL-70, RNA-Pol-III |
| 81 | F | 2019_03_22 | 78 | Skin, Heart | 25 | Steroids >12,5mg | RNA-Pol-III |
| 58 | M | 2019_03_27 | 55 | Skin | 19 | MMT | SCL70 |
| 59 | M | 2021_03_21 | 55 | Skin, Lungs | 20 | MMT | SCL70 |
| 66 | M | 2019_04_01 | 65 | Skin, Heart, Lungs, Kidneys, Digestive | 27 | Aspirin | SCL70 |
| 29 | F | 2019_04_15 | 27 | Skin, Heart, Lungs, Digestive | 21 | None | SCL70 |
| 44 | M | 2019_04_30 | 44 | Skin, Digestive | 14 | None | SCL70 |
| 29 | M | 2019_06_12 | 23 | Skin | 12 | None | SCL70 |
| 35 | F | 2019_07_11 | 34 | Skin, Lungs, Digestive | 25 | MMT | SCL70 |
| 46 | M | 2021_12_08 | 45 | Skin, Lungs, Digestive | 27 | MMT | SCL70, Centromere |
| 36 | F | 2022_01_26 | 32 | Skin, Lungs | 16 | MMT | SCL70 |
| 18 | M | 2022_06_08 | 17 | Skin, Lungs, Digestive, Articular | 26 | MMT, Steroids >12,5mg | SCL70 |
| 25 | F | 2022_06_22 | 23 | Skin | 20 | None | SCL70 |
| 55 | M | 2022_09_14 | 55 | Skin | 18 | MMT, Steroids >12,5mg | nuclear |
| 47 | F | 2021_08_07 | 45 | Skin | 19 | None | SCL70 |
| 25 | F | 2023_02_17 | 20 | Skin, Lungs, Digestive | 26 | MMT | SCL70 |
| 41 | F | 2023_03_08 | 32 | Skin, Lungs, Digestive | 29 | None | SCL70 |
| 47 | F | 2023_03_10 | 34 | Skin, Lungs, Digestive | 32 | None | SCL70 |
| 60 | F | 2023_06_19 | 59 | Skin, Lungs | 18 | MMT | SCL70 |
| 58 | F | 2023_06_19 | NC | Skin, Lungs, Digestive | 21 | MMT | SCL70 |

**Table 2:** Systemic lupus erythematosus patients informations. Age, sex, date of prelevement, age of first sign, clinical manifestations, SLEDAI score, treatments at the time of prelevement and presence of auto antibodies. NSAID : Non-steroidal anti-inflammatory drugs

| Age | Sex | Prelevement date | Age of first sign | Clinical manifestations | SLEDAI score | Treatments | Auto-antibodies |
| --- | --- | --- | --- | --- | --- | --- | --- |
| 43 | F | 2022_02_15 | 25 | Skin, articular, pericarditis | 0 | Hydroxychloroquine | dsDNA, anti-nucleosome, anti-SSA |
| 42 | F | 2023_03_02 | 36 | Skin | 2 | Steroids 5mg | dsDNA, anti-SSA |
| 47 | F | 2023_03_02 | 43 | Skin, acticular | 8 | NSAID | dsDNA, anti-SSA |
| 54 | F | 2023_06_19 | 43 | Skin, articular | 2 | Hydroxychloroquine, Steroids 20mg | dsDNA |

### PBMC isolation from blood

Blood is diluted in preheated PBS (1:1 ratio) and added on top of a Ficoll layer (2:1 ratio) and centrifuged at 1200*g for 20 minutes with acceleration and brakes set to a minimum. PBMC are harvested and washed with PBS. Platelets are removed by 2 centrifugations at 120*g for 10 minutes with brake and acceleration set to low and then counted.

### T cell Isolation and *in vitro* stimulation

T cells are isolated from freshly harvested PBMC using human Pan T cell Isolation following manufacturer instructions. 2*10^5^ cells are seeded per well of 96 multi well plates and stimulated overnight with 50ng/ml phorbol myristate and 1μM ionomycin or left unstimulated.

### *In vitro* monocyte-derived macrophage differentiation and phenotyping

Monocytes are purified from freshly isolated PBMC by CD14^+^ positive magnetic sorting following manufacturer instructions. Monocytes are then seeded in multi well plates at 1*10^5^ cells/cm^2^ and incubated for 3 days at 37°C 5% CO_2_ in presence 50ng/mL of M-CSF for M0 and M2 differentiation and 10ng/mL of GM-CSF for M1 differentiation. On day 3, cytokines are renewed. On day 6, 10ng/mL of lipopolysaccharide and 50ng/mL of IFN-γ are added to monocytes for M1 differentiation and 20ng/mL of IL-4 are added to monocytes for M2 differentiation. On day 7, macrophages are harvested with accutase and stained for extracellular markers (anti CD163-AF647, anti CD206-PerCP-eFluor 710, anti CD282-AF488, anti CD14-BV510, anti HLA-DR-BV711, anti-CD16-APC/Cyanine7, anti-CD80-PE/Cyanine7, anti CD86-PE/CF594, anti-CD369-PE) for phenotyping by flow cytometry or seeded in 8 chamber slides for phagocytosis assays. Flow cytometric data are acquired with an Attune NxT flow cytometer (Thermo Fisher Scientific) and data analyzed with FlowJo V10 (LLC).

### Quantification of LC3 accumulation by flow cytometry

1*10^5^ cells are seeded per well of 96 multi well plates. Cells are treated with Autophagy Reagent A or left unstimulated for 2 hours at 37°C 5% CO_2_ in complete medium: RPMI 1640 supplemented with 10% (v/v) heat decomplemented fetal bovine serum and 50μg/mL of gentamycin. Cells are stained with anti CD3-APC-AF750, anti CD19-APC and anti CD14-PerCP-Cy5.5 and then permeabilized with 0.05% (m/v) saponin. Soluble LC3 is washed out by centrifugation at 500*g for 5 minutes and cells are then stained with anti-LC3-FITC antibody. Acquisition is done in assay buffer with a Gallios flow cytometer (Beckman Coulter), and data analyzed with FlowJo V10 (LLC). Autophagic flux is calculated by dividing the LC3-FITC mean fluorescence intensity (MFI) of the Autophagy Reagent A treated condition by the MFI of the untreated condition.

### Phagocytosis assays and anti-LC3B immunofluorescence

100μg of Human Immunoglobulin type G (Sigma-Aldrich, 14506) or Bovin Serum Albumin are covalently linked to 3μm Polybead® Carboxylate Microspheres with PolyLink Protein Coupling Kit following manufacturer instructions. 2*10^5^ macrophages are seeded per well of 8 chamber slides and left to attach overnight. 8*10^5^ beads are incubated with macrophages for 45 minutes at 37°C 5% CO_2_ in complete medium. Cells are then washed twice with ice cold PBS to remove free beads. Cells are fixed with 4% (v/v) paraformaldehyde diluted in PBS for 10 minutes at room temperature and washed 3 times with PBS. Cells are then permeabilized by 30 minutes incubation with 0.05% (v/v) Triton X-100 diluted in PBS 2% BSA (v/m). Blocking is then done by incubation with PBS-2% BSA (v/m) for 1 hour. Primary anti-LC3B antibody diluted in PBS 2% BSA is then incubated overnight at 4°C. Cells are washed 5 times with PBS and incubated 1 hour at room temperature with secondary goat anti-rabbit IgG Alexa Fluor™ Plus 647 antibodies diluted in PBS 2% BSA. Cells are washed 5 times for 5 minutes with PBS and mounted with ProLong™ Glass Antifade Mountant with NucBlue™. After overnight curing, slides are randomly imaged with a Zeiss LSM 800 AiryScan microscope with 40x/1,4 (DIC) NA objective. 1μm z-sections are acquired and phagosomes and LAPosomes are manually counted using Fiji (https://imagej.net/).

### Statistics

Data are reported as means ± standard deviation and analyzed by with two-tailed Wilcoxon test, two ways ANOVA with Tukey multiple comparison or Kruskal-Wallis test with Dunn’s multiple comparisons test using GraphPad Prism 8 (GraphPad Software) or BioRender (Biorender.com). All statistical analysis was considered significant at P < 0.05.

Antibody and reagents

Antibody and reagents.
| Reagent | Concentration or dilution | Reference |
| --- | --- | --- |
| **8 Chamber slides** | - | Milipore, PEZGS0816 |
| **PBS** | - | Sigma-Aldrich, D8537 |
| **Ficoll Paque PLUS** | - | Sigma-Aldrich, 17-1440-03 |
| **RPMI 1640** | - | Gibco™, 21875034 |
| **Fetal bovine serum** | 10% (v/v) | Gibco™, A3160802 |
| **Gentamycin** | 50µg/mL | Gibco™, 15750037 |
| **Accutase** | - | Gibco™, A1110501 |
| **Pan T cell Isolation Kit, human** | - | Miltenyi Biotec, 130-096-535 |
| **CD14 MicroBeads, human** | 20µL/10<sup>7</sup> cells | Miltenyi Biotec, 130-050-201 |
| **Autophagy Reagent A** | 1/1000 | Luminex®, FCCH10017 |
| **Phorbol myristate acetate** | 50ng/mL | Invivogen, tlr-pma |
| **Ionomycin** | 1µM | Invivogen, inh-ion-3 |
| **M-CSF** | 50ng/mL | R&D, 216-MC |
| **GM-CSF** | 10ng/mL | Gift from F. Oberling (CHUS) |
| **Lipopolysaccharide** | 10ng/mL | Sigma-Aldrich, 297-473-0 |
| **IFN-γ** | 50ng/mL | Peprtech, 200-04 |
| **IL-4** | 20ng/mL | Peprtech, 200-04 |
| **Saponin** | 0,05% (m/v) | Sigma-Aldrich, S4521 |
| **Triton X-100** | 0,05% (v/v) | Euromedex, 2000-C |
| **Paraformaldehyde** | 4% (v/v) | EMS, 15710 |
| **ProLong™ Glass** | - | Invitrogen, P36985 |
| **Human Immunoglobulin type G** | - | Sigma-Aldrich, 14506 |
| **Bovin Serum Albumin** | - | Euromedex, 04-100-812-E |
| **3µm Carboxylate Microspheres** | - | Polysciences, 09850-5 |
| **Protein Coupling Kit** | - | Polysciences, 24350-1 |
| **Anti CD3-APC-AF750 (UCHT1)** | FC: 1/100 | Beckman Coulter, A94680 |
| **Anti CD19-APC (HIB19)** | FC: 1/100 | BD Bioscience, 555415 |
| **Anti CD14-PerCP-Cy5.5 (M5E2)** | FC: 1/100 | BD Bioscience, 550787 |
| **Anti LC3-FITC (4E12)** | FC: 1/20 | Luminex®, FCCH10017 |
| **Anti LC3 (polyclonal)** | IF: 1/200 | Abcam, ab48394 |
| **Anti CD163-AF647 (MAC-2-158)** | FC: 1/400 | Invitrogen, 51-1637-42 |
| **Anti CD282-AF488 (TL2.1)** | FC: 1/100 | Luminex®, FCCH10017 |
| **Anti CD206-PerCP-eFluor 710 (19.2)** | FC: 1/100 | Invitrogen, 46-2069-42, |
| **Anti CD14-BV510 (MφP-9)** | FC: 1/200 | BD Biosciences, 563079 |
| **Anti HLA-DR-BV711 (G46-6)** | FC: 1/500 | BD Biosciences, 563696 |
| **Anti CD16-APC-Cy7 (3G8)** | FC: 1/500 | BD Biosciences, 302018 |
| **Anti CD80 -PE-Cy7 (L307,4)** | FC : 1/400 | BD Biosciences, 561135 |
| **Anti CD86-PE-CF594 (2331)** | FC : 1/500 | BD Horizon, 562390 |
| **Anti CD369-PE (15E2)** | FC : 1/100 | Invitrogen, 12-9856-42 |
| **Anti Rabbit-IgG-AF+647 (polyclonal)** | IF: 1/500 | Invitrogen, P36985 |

## Results Discussion

Autophagy could contribute to autoreactive lymphocyte survival. As we and others previously described autophagy dysregulations in T and B lymphocytes from systemic autoimmune diseases, we investigated by flow cytometry the levels of LC3B-II in peripheral blood mononuclear cells (PBMC) from SSc patients and age and sex matched healthy controls. All patients fulfilled 2013 ACR/EULAR classification criteria^15^ (table I). Briefly, cells are treated with saponin without prior fixation, to eliminate soluble cytosolic LC3-I and stain only LC3-II that is associated with membranes. This staining method allows to assess in each sample LC3-II levels in B and T lymphocytes as well as monocytes (Figure 1A and B). We did not detect significant differences in LC3-II at steady state between healthy donors (HD) and SSc patients, in T cells, B cells nor monocytes (Figure 1C). LC3-II can be used as a marker for autophagosomes but is also associated with other intracellular vesicles during CASM. To focus on autophagic activity, we then assessed the autophagic flux by using bafilomycin A1 (BafA1) that blocks autophagosome degradation. As LC3-II is continuously recycled at the final steps of autophagy, this treatment leads to accumulation of LC3-II positive autophagosomes. Although autophagic flux was detectable in T cells and monocytes, no difference in LC3-II accumulation could be noticed between HD and SSc macrophages (Figure 1D). We could however observe a slight reproducible flux increase in SSc B cells compared to HD (5 patients out of 7 tested). As it was previously described that T cell activation leads to accumulation of LC3-II, we checked in PMA-Ionomycin stimulated cells if differences could be observed in these conditions. Consistent with results obtained in fresh PBMCs, unstimulated T cells show no difference LC3-II expression level, or flux between HD and SSc samples. We report however a slight increase LC3-II level at basal and decrease after BafA1 stimulation. This could reflect a blockade in the autophagic process at the degradation step in SSc T cells, as shown by the flux calculation (Figure 1D).

**Figure 1:**
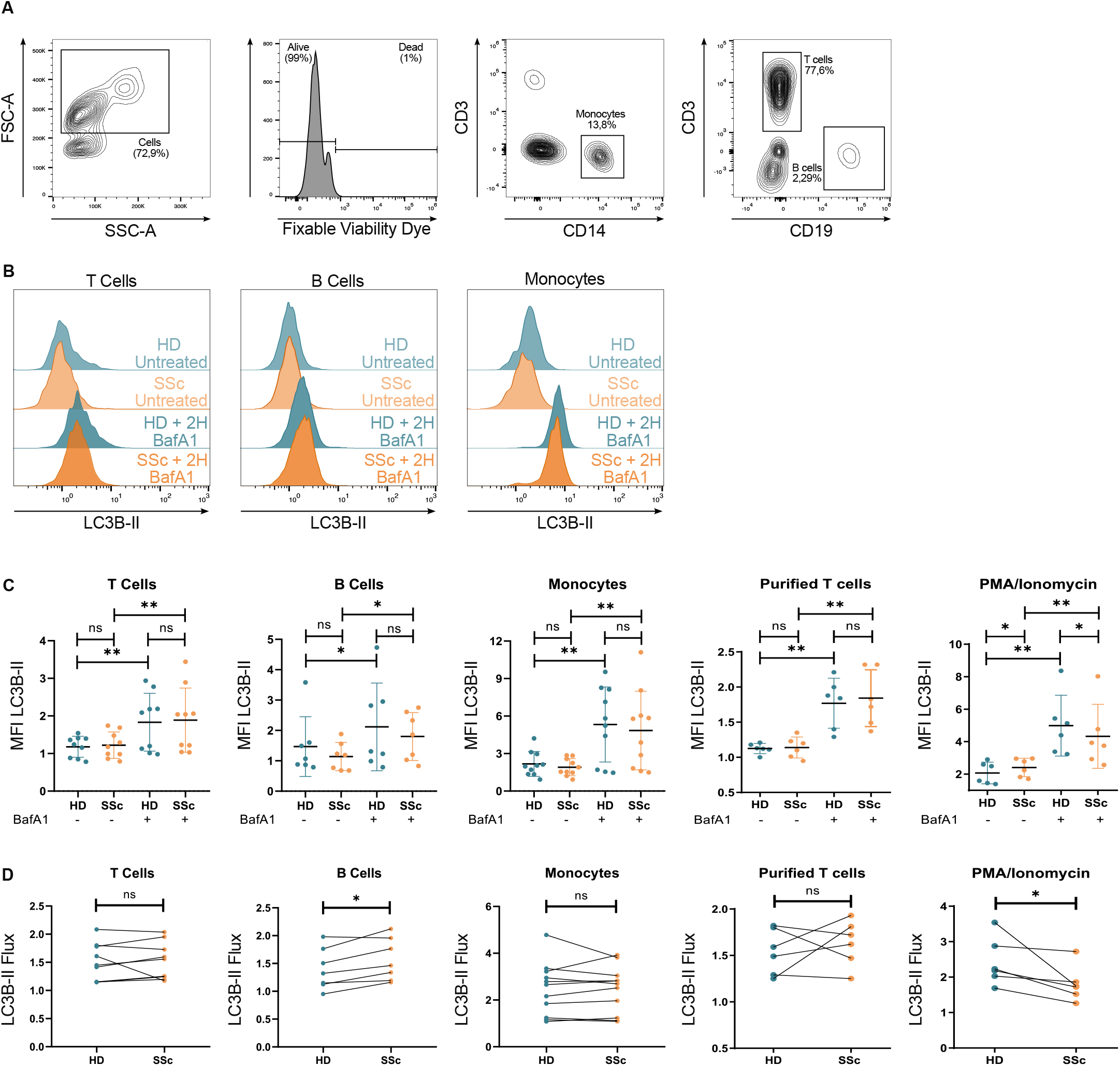
Autophagic flux is not different in PBMC from SSc patients. Gating strategy (A). Histogram plot of LC3-II MFI after 2 hours Bafilomycin A1 treatment in T cells, B cells and monocytes (B). LC3-II MFI after 2 hours of Bafilomycin A1 treatment in T cells, B cells, monocytes, untreated purified T cells or purified T cells treated with PMA/ionomycin (C). LC3-II Flux in T cells, B cells, monocytes, untreated purified T cells or purified T cells treated with PMA/ionomycin (D). Statistical analyses are performed with two-tailed Wilcoxon tests (not significant=ns;*=p<0.05; **=p<0.005; ***=p<0.001; ****=p<0,0001).

We then decided to investigate Atg8ylation in monocyte-differentiated macrophages. To do so, we differentiated macrophages from purified circulating CD14^+^ monocytes. As Atg8ylation processes could be regulated differently according to macrophage polarization, we differentiated macrophages from purified circulating monocytes under M0, M1 or M2 skewing conditions (Figure 2A). We also included samples from another systemic autoimmune disease, SLE, for comparison. We could find preferential expression of the expected markers in the corresponding polarization settings for HD, as well as SSc and SLE samples (HLA-DR, CD80, CD86 for M1 cells, CD163, CD16 for M0 and M2 cells and Dectin 1 for M2 cells). Interestingly, we found a tendency for decreased HLA-DR and CD86 expression in SSc and SLE M1 samples (Figure 2B). We also found that for SSc and SLE patients, a high number of cells were both CD163 and CD80 under M1 skewing condition indicating a potential bias toward and intermediate M1/M2 phenotype (Figure 2C, 2D). This finding is consistent with previous reports mentioning a propensity for M2 or M2-like macrophage differentiation, or mixed M1/M2 phenotypes in SSc tissues^16^. We found no major difference in M0 or M2-associated marker expression between SSc, SLE patients, and HD macrophages. We then assessed LC3-II flux in M0 macrophages from SSc patients in four patients. As for non-differentiated monocytes, we found no tendency for dysregulated autophagy in these cells (Figure 2E).

**Figure 2:**
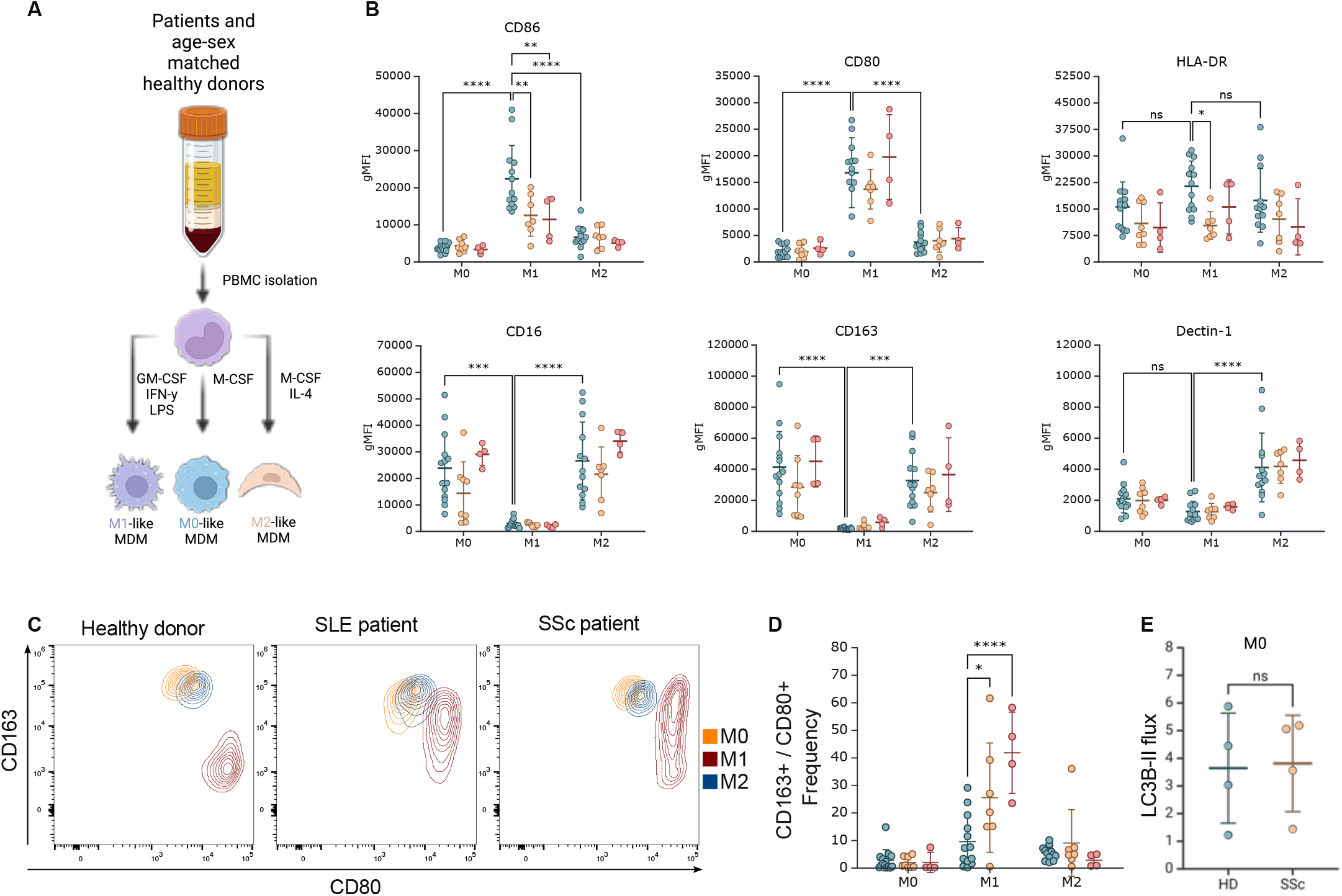
M1 but not M0 or M2 macrophages differenciation is impaired in patients compared to healthy donors. CD14-positive monocytes were isolated from peripheral blood mononuclear cells and differentiated with Granulocyte-Macrophage Colony Stimulating Factor (10ng/mL), Interferon-gamma (50ng/mL), Lipopolysaccharide (10ng/mL) for M1-like monocytes-derived macrophages (MDM) ; Macrophage Colony Stimulating Factor (50ng/mL) for M0-like MDM ; Macrophage Colony Stimulating Factor (50ng/mL), Interleukine-4 (50ng/mL) for M2-like MDM (A). Geographic mean fluorescence intensity of CD86, CD80, HLA-DR, CD16, CD163 and Dectin-1 (B) analyzed by flow cytometry. Statistical analyses are performed with two ways ANOVA with Tukey multiple comparison (*=p<0.05; **=p<0.005, ***=p<0.001, ****=p<0,0001). Dot plot (C) and associated proportion of CD163/CD80 double positive populations (D) under M0, M1 or M2 skewing conditions in healthy donor, SLE patient or SSc patient. LC3B flux in M0 macrophages quantified by flow cytometry (E). Statistical analyses are performed with two ways ANOVA with Tukey multiple comparison for B and D or with two-tailed Wilcoxon test for E (not significant=ns;*=p<0.05; **=p<0.005; ***=p<0.001; ****=p<0,0001).

As differentiation of M0 cells and autophagy levels are comparable between SSc patients and HD, we decided to investigate if other Atg8ylation processes active in professional phagocytes could be impaired. To evaluate LAP activity, we incubated macrophages with beads coated with LAP-inducing stimulus (IgG) and non-coated beads that should only engage phagocytosis (Figure 3A). We could first show the specific induction of LAP with IgG-coated beads, compared with non-coated beads than only induce phagocytosis (Figure 3B). We then compared between patients and controls the level of phagosomes per cell of SSc macrophages compared to controls. We found no significant difference between SSc and HD (Figure 3C). Interestingly, SLE samples tested were in the low interval, consistent with phagocytic defects characteristic of the pathology. To measure the level of LAP, we stained the samples with anti-LC3 antibody, allowing to discriminate phagosomes from LAPosomes (LC3-II+ phagosomes, Figure 3B). We found a reproducible decrease of the number of LAPosomes in SSc macrophages versus their matched control (Figure 3D). This tendency was confirmed by calculating the ratio of laposomes per total phagocytic vesicles, where we found a strong decrease of LAP activation (Figure 3E). In contrast, the SLE samples tested were in a higher range of LAPosomes frequency found in SSc, but still in the low range compared with HD.

**Figure 3:**
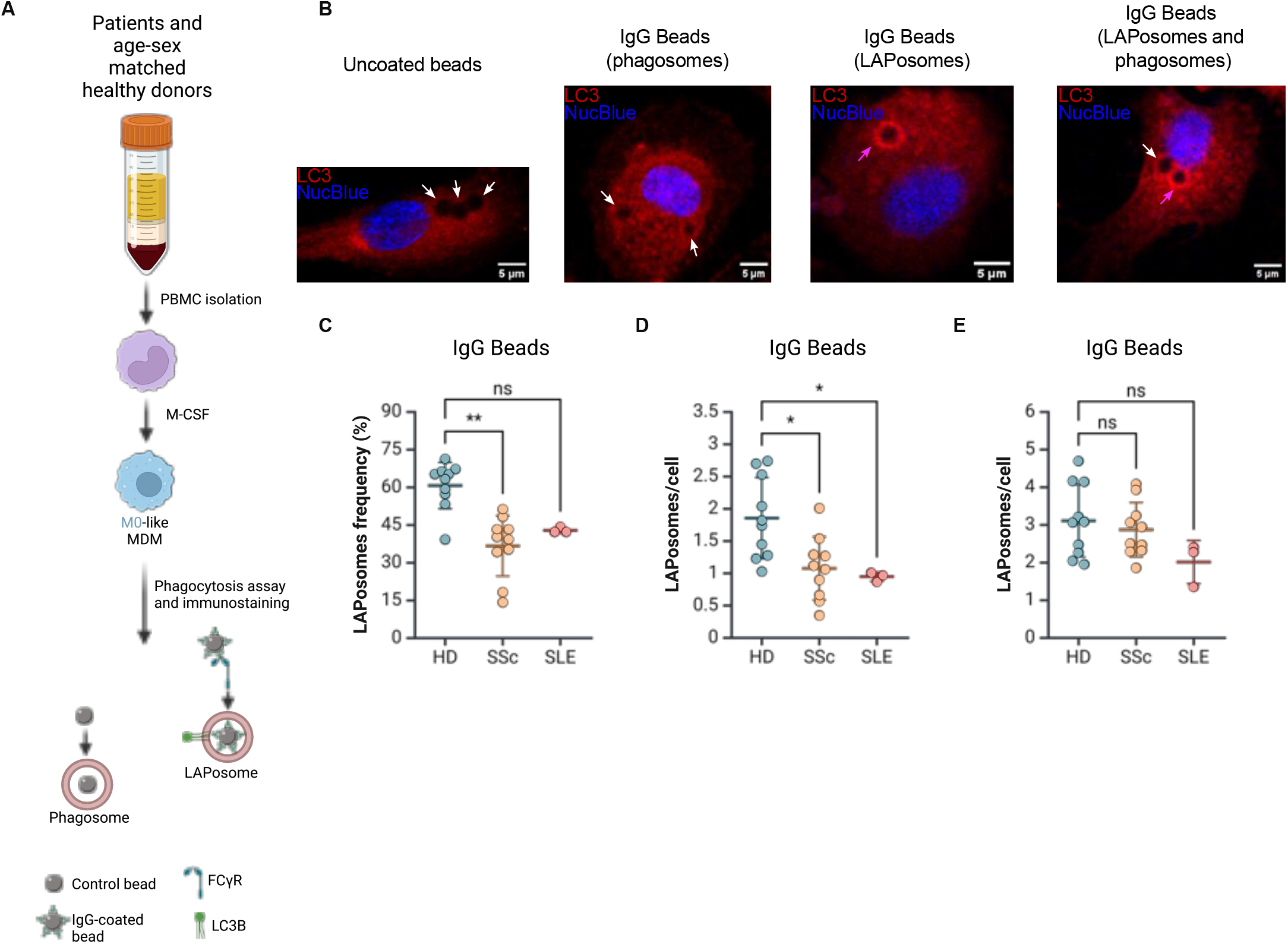
LC3-associated phagocytosis is defective in M0-like monocytes-derived macrophages from Systemic sclerosis patients. CD14-positive monocytes were isolated from peripheral blood mononuclear cells and differentiated with Macrophage Colony Stimulating Factor (50ng/ mL) for M0-like monocytes-derived macrophages (MDM). Phagocytosis assays with Immunoglobulin type-G (IgG) were performed on 8-chambers slides for 45 minutes, fixed and immunostained with anti-LC3B antibody. LAPosomes are specifically induced after IgG recognition by Fc-gamma receptor (FcgR) (A). Representative images of anti-LC3 immunofluorescence with cells with phagosomes (white arrow) and/or LAPosomes (magenta arrows) (B). Mean number of phagosomes/cell (C), LAPosome/cell (D) and LAPosome frequency (E), and after phagocytosis of IgG coated beads. Each dot represents an independent experiment with at least 45 cells. Statistical analyses are performed with Kruskal-Wallis test with Dunn’s multiple comparisons test (not significant=ns;*=p<0.05; **=p<0.005; ***=p<0.001; ****=p<0,0001).

We thus show that no deregulation of autophagy is found in SSc circulating lymphocytes and monocytes. We however find in cultured T cells, a tendency for increased LC3-II staining, and less accumulation after BafA1 treatment. This is reminiscent of what is found for SLE T cells in several reports, showing abnormal accumulation of autophagic vesicles^17^ or LC3-positive compartments^18,19^. What remains unclear is if it is the consequence of more autophagosome generated, less lysosomal degradation or both. In this study, we find a decreased autophagic flux that can be the consequence of both phenomena. The most striking results we report is the intrinsic impairment of LAP in SSc and SLE macrophages. This seems more pronounced in SSc although more SLE samples would be needed to draw a definitive conclusion for this comparison. Interestingly, LAP has been shown to be induced in liver fibrosis and protective against inflammation^20^. Inhibition of LAP exacerbated inflammatory signature of human monocytes from cirrhotic patients and from mouse models with chronic liver injury. LAP is proposed to be induced in this context by engagement of immune complexes by FcγRIIA, leading to inhibitory ITAM (ITAMi) signaling and recruitment of the phosphatase SHP-1 which could explain anti-inflammatory effects. In SSc we can speculate that defects in Fc receptor signaling are involved in the impaired LAP activation. Interestingly, CD16 expression is linked to the M2 phenotype, that is the most efficient in efferocytosis. We can thus speculate that LAP is more active in M2 macrophages. In the context of SSc, in M0 cells where we assessed LAP activity, no significant difference could be found in CD16 surface expression between SSc and HD. This means that intrinsic signaling deregulation are more likely responsible for LAP impairment. It would be interesting to investigate FcγR downstream signaling such as SYK recruitment, ITAMi activation and association with SHP-1.The ATG16L1/V-ATPase axis is central to Atg8ylation in the context of LAP and more generally, CASM^21^. It would be also interesting to test if Atg8ylation occurs in response to STING activation that is exacerbated in SSc^22^. The consequences of LAP defects in SSc still remain to be defined. It was proposed that efferocytic defects in SSc participate to accumulation of cell debris and related inflammation and autoantigen presentation as it in the case in SLE^4,5^. Several reports, including ours, show that monocyte-derived macrophages are skewed in certain conditions toward atypic mixed M1/M2 phenotypes^16^. It would be interesting to systemically evaluate LAP activity in known macrophage polarized phenotypes and compare it to global efferocytic and phagocytic capacities.

Efferocytosis defects are central in SLE. Interestingly, several lines of evidence point towards shared physiopathological features between, SLE and SSc. Efferocytic defects could indeed lead in both pathologies to IFN signatures observed in several SSc patients^23^. Interestingly, LAP has been involved in the limitation of IFN signaling after engulfment of dead cells^12^. LAP defects in SSc could contribute to hyperinflammation and IFN signature observed in SSc patients.

Our study points towards a possible pathological mechanism linked to LAP activity in macrophages. We need now to evaluate the impact of LAP on inflammatory potential of SSc macrophages. Specific LAP modulators could be envisaged as therapeutic options^10,24^. It was recently shown that columbamine enhance macrophage-mediated efferocytosis by promoting LAP to protect intestinal inflammation in a murine colitis model^25^. LAP activity could attenuate fibrosis in SSc as proposed by others. We propose that it could also attenuate autoimmune and inflammatory responses.

## Contributorship

F.G and T.M.designed the research.

F.G, Q.F, J.L., A.P, L.P. performed the experiments.

F.G, Q.F, J.L., A.P, L.P analyzed the data.

F.G, Q.F, J.L wrote the paper.

## Abbreviations

ATG: (autophagy related)
CASM: (conjugation of ATG8 to single membranes)
IFN: (interferon)
LAP: (LC3-associated phagocytosis)
MAP1LC3B or LC3B: (microtubule-associated protein light chain 3B or light chain 3B)
SLE: (systemic lupus erythematosus)
SSc: (systemic sclerosis)
PBMC: (peripheral blood mononuclear cells)
TLR: (toll-like receptor)

## Acknowledgements

We thank Valérie Lecureur and Alain Lescoat for valuable discussions and exchanges.

We thank Pascal Kessler and the PICSTRA platform (Centre de Recherche en Biomédecine de Strasbourg, UMS38 INSERM) for technical help.

## Ethical Approvements

This study was approved and done in the frame of RESO, national reference center for rare diseases. All patients signed a consent form after being informed about the protocol and the aim of this study.

## Data sharing

Authors agree to share any relevant data upon request

## Patient and Public Involvement

Non applicable

